# Identification of a fatty acid-sensitive interaction between mitochondrial uncoupling protein 3 and enoyl-CoA hydratase 1 in skeletal muscle

**DOI:** 10.1101/821637

**Authors:** Christine K. Dao, Alexander Kenaston, Katsuya Hirasaka, Shohei Kohno, Christopher Riley, Gloria Fang, Kristin Fathe, Ashley Solmonson, Sara M. Nowinski, Matthew E. Pfeiffer, Xianmei Yang, Takeshi Nikawa, Edward M. Mills

## Abstract

Skeletal muscle mitochondrial fatty acid (FA) overload in response to chronic overnutrition is a prominent pathophysiological mechanism in obesity-induced metabolic disease. Increased disposal of FAs is therefore an attractive strategy for intervening in obesity and related disorders. Skeletal muscle uncoupling protein 3 (UCP3) activity is associated with increased FA oxidation and antagonizes weight gain in mice on obesogenic diets, but the mechanisms involved are not clear. Here, we show that UCP3 forms a direct, FA-stimulated, mitochondrial matrix-localized complex with the auxiliary unsaturated FA-metabolizing enzyme, Δ^3,5-^Δ^2,4^dienoyl-CoA-isomerase (ECH1). Expression studies in C2C12 myoblasts that functionally augments state 4 (uncoupled) respiration and FA oxidation in skeletal myocytes.

Mechanistic studies indicate that ECH1:UCP3 complex formation is likely stimulated by FA import into the mitochondria to enhance uncoupled respiration and unsaturated FA oxidation in mouse skeletal myocytes. In order to characterize the contribution of ECH1-dependent FA metabolism in NST, we generated an ECH1 knockout mouse and found that these mice were severely cold intolerant, despite an up-regulation of UCP3 expression in SKM. These findings illuminate a novel mechanism that links unsaturated FA metabolism with mitochondrial uncoupling and non-shivering thermogenesis in SKM.

## Introduction

The accumulation of fatty acids (FAs) in skeletal muscle (SKM), caused by chronic over-nutrition where the energetic supply exceeds mitochondrial oxidative capacity, is a prominent patho-physiological mechanism linking obesity to insulin resistance and related metabolic diseases (Krssak et al. 1998; Pan et al. 1997; Roden et al. 1996; Boden 2011). Mitochondrial dysfunction is common in obese patients and is associated with defects in fatty acid (FA) disposal (Lowell & Shulman 2005; Morino et al. 2006). Thus, strategies to dissipate the energy surplus and increase FA metabolism to relieve mitochondrial overload are promising therapeutic approaches for combating metabolic disease. Mitochondrial uncoupling proteins (UCPs) comprise a subfamily of the mitochondrial solute carrier (Slc) superfamily that uncouple the mitochondrial respiratory chain from ATP synthesis through the regulation of inner mitochondrial membrane proton leak (Krauss et al. 2005) in a variety of tissues, including brown adipose tissue (BAT), heart, SKM (Cannon et al. 1982; Brand & Esteves 2005), and most recently skin (Lago et al. 2012). The canonical UCP homologue, UCP1, is required for adaptive non-shivering thermogenesis (NST) in BAT-the mitochondrial bioenergetic process by which the energy stored in fuel substrates are rapidly metabolized and released in the form of heat in response to cold (Enerback et al. 1997).

Although the UCP1 homologue, UCP3, is not recognized as a direct regulator of adaptive NST, there is considerable evidence to support the role of UCP3 as a physiological mediator of energy balance and fatty acid metabolism (Samec et al. 1998; Brand & Esteves 2005). The induction of UCP3 expression in physiological states where FA levels are high, such as high fat feeding and fasting, suggests that UCP3 is important in mediating lipid handling and preventing mitochondrial overload (Bézaire et al. 2001). Similarly, clinical studies have linked genetic mutations that alter UCP3 function to the development of obesity and type II diabetes in susceptible human populations (Schrauwen et al. 1999; Schrauwen et al. 2001; Liu et al. 2005; Musa et al. 2011). In addition to this, animal studies have shown that overexpression of UCP3 in SKM not only increases FA metabolism and protects against oxidative stress (MacLellan et al. 2005), but also enhances glucose homeostasis (Choi et al. 2007; Huppertz et al. 2001; Clapham et al. 2000). However, the molecular mechanisms that govern UCP3 function in obesity and FA metabolism are not well understood.

This study advances our understanding of the mechanisms that link UCP3 function to FA metabolism in SKM, as a means to relieve mitochondrial overload and metabolic stress. Here we show that UCP3 directly interacts with the auxiliary, unsaturated FA metabolizing enzyme enoyl CoA hydratase-1 (ECH1) in the mitochondrial matrix. Unlike saturated FAs that can readily undergo mitochondrial β-oxidation, unsaturated FA metabolism is more complex and requires a set of specific auxiliary enzymes that permit the complete oxidation of these particular FA species. ECH1 is an essential enzyme in the reductase-dependent pathway, which is one of the two branches that can metabolize unsaturated FAs with double bonds in odd-numbered positions along the carbon chain (Luo et al. 1994). In the reductase-pathway, ECH1 catalyzes the isomerization of a 3-trans, 5–cis dienoyl-CoA substrate to a 2-trans, 4-trans dienoyl-CoA product (Filppula 1998; Luthria et al. 1995)-a step that permits the complete oxidation of these specific FA metabolites. Although little is known about the physiological relevance of ECH1, it has been proposed that flux through the reductase-dependent pathway is important in facilitating complete FA metabolism, thus protecting mitochondrial oxidative function and preventing metabolic stress (Shoukry & Schulz 1998). This hypothesis is supported by studies where accumulation of the 3,5 enoyl-CoA derivative is shown to strongly inhibit beta-oxidation in *E. coli* that do not endogenously express ECH1 (Ren et al. 2004). Additionally, FA levels are significantly elevated following ECH1-knockdown in *C. elegans* compared to wild type, suggesting that normal β-oxidation is compromised in this model. Furthermore, mice lacking dienoyl-CoA reductase, an enzyme that acts directly downstream of ECH1 and is also involved in the metabolism of unsaturated FAs with even-numbered double bonds, showed a compromised thermoregulatory response when challenged by fasting and cold exposure. Together, these studies indicate that the auxiliary enzymes involved in the complete breakdown of unsaturated fatty acids are critical in the adaptation to metabolic stress (e.g. fasting) by maintaining balanced FA and energy metabolism.

Our work demonstrates that ECH1 and UCP3 form a novel protein-protein interacting complex that is regulated by FAs and likely important in the adaptation to metabolic stress. UCP3 and ECH1 directly interact at endogenous concentrations, and the presence of both proteins enhances uncoupled respiration and unsaturated FA oxidation. To further characterize the importance of the synergistic relationship between ECH1 and UCP3 in SKM metabolism, we generated two ECH1-knockout mouse lines, and found that these mice were unable to defend their core body temperature when challenged with fasting and cold-exposure, despite a compensatory increase in UCP3 protein expression. To our knowledge this is the first demonstration of a protein-protein interaction formed in the mitochondrial matrix with any UCP protein.

## Materials and Methods

### Chemicals and Reagents

Unless otherwise noted, all reagents were purchased from Sigma (St. Louis, MO).

### Plasmid DNA constructs

Full-length UCP3, UCP1 and ECH1 genes were amplified from mouse heart cDNA library and cloned into pcDNA3.1 (Invitrogen, Carlsbad, CA) with either the C-terminal V5 or Myc tag, respectively. Site-directed mutagenesis of plasmids to generate the C-terminal Myc tagged ECH1 catalytic mutants and the C-terminal V5 tagged UCP3 truncation mutants was carried out with Platinum pFX DNA polymerase from (Life Technologies, Grand Island, NY), following the manufacturer’s protocols for site-directed mutagenesis. The primers used to generate the Myc tagged-ECH1 catalytic mutants were previously published in Zhang *et al*.

### Cell culture

HEK293T and C2C12 cells were obtained from the American Type Culture Collection (ATCC, Manassas, VA). HEK293T cells were cultured at 37°C with 5% CO_2_ in Dulbecco’s Modified Eagle Medium (Cellgro, Manassas, VA) containing 10% Fetal Bovine Serum1% and 100X PenG-Streptomycin (Invitrogen, Carlsbad, CA). C2C12s were cultured in similar conditions, with the exception of being grown in high glucose (45000mg/L), Dulbecco’s Modified Eagle Medium from Sigma (St. Louis, MO).

### Transient transfection of cells

HEK293Ts were plated one day prior to transfection and were transfected with either calcium phosphate or TransIT-LT1 (Mirus, Madison, WI) following the manufacturer’s instructions. To improve transfection efficiency of C2C12s, a cell line that is generally difficult to transfect, cells were transfected with Lipofectamine 2000 at a 2:1 ratio (Invitrogen, Carlsbad, CA) immediately after plating cells, according to Mercer *et al*.

### Generation of stable C2C12 cell lines

The Precision LentiORF lentiviral packaging system was obtained from Thermo Fisher Scientific (Waltham, MA). Full-length ECH1 was cloned into the pLOC lentiviral plasmid following the manufacturer’s instructions. HEK293T packaging cells were transfected with the pLOC lentiviral plasmid and two viral packaging plasmids pMDG2 and psPAX2, using a standard calcium phosphate transfection method to produce the lentiviral particles. The HEK293T cells were incubated with the transfection complexes in normal growth media supplemented with 25μM choloroquinone for 8 hours. Forty hours after transfection, the virus containing media was harvested. To concentrate the lentiviral particles, the Lenti-X Concentrator reagent (Clonetech, Mountainview, CA) was used according to the manufacturer’s protocols. For lentiviral transduction, C2C12 myocytes were seeded in 6 well plates at 50,000 cells/wells one day prior to infection. The cells were then incubated with the concentrated lentiviral particles in full growth media supplemented with polybrene (8μg/mL), overnight. Following infection, stable cell colonies were selected with 25μg/mL of blasticidin. Positive colonies were then confirmed through immunoblotting and densitometry.

### Animals

C57BL/6J wild-type mice were obtained from Jackson Laboratories (Bar Harbor, ME). The UCP3 knockout mice (UCP3^-/-^) in the C57BL/6J background were a gift from Dr. Marc Reitman, formerly of the National Institutes of Health. The UCP1 knockout mice (UCP1^-/-^) were a gift from Dr. Leslie Kozak of the Pennington Biomedical Research Institute. The UCP3 ^-/-^ mice were crossed with the UCP1^-/-^ mice to generate our UCP1 and UCP3 double knockout line (UCP1^-/-^ + UCP3^-/-^). The transgenic mice overexpressing human UCP3 in SKM generated from human alpha1-actin promoter targeting construct (TgSKM UCP3 ^+/+^) were a gift from Dr. Mary-Ellen Harper of the University of Ottawa. The TgSKM UCP3^+/+^ were then crossed with the UCP3 -/- and DKO lines to generate a transgenic mouse overexpressing human UCP3 in SKM in the UCP3^-/-^ and DKO background (TgSKM UCP3 ^-/-^ and TgSKM UCP1^-/-^ + UCP3^-/-^, respectively). It is important to note that all TgSKM-UCP3 strains were kept as hemizygous breeder colonies.

Unless otherwise noted, all experiments were performed in male mice between the ages of 6-8.5 weeks. All animal husbandry and procedures were carried out in accordance to the Association for Assessment and Accreditation of Laboratory Animal Care (AAALAC) and approved by the Institutional Animal Care and Use Committee at The University of Texas at Austin (IACUC).

### Generation of ECH1 ^-/-^ mouse model

The custom designed CompoZr^®^ Zinc Finger Nuclease Plasmid targeted to exon 3 of the mouse ECH1 gene on chromosome 7 was obtained from Sigma. In-vitro transcription of the ZFN mRNA was performed following the manufacturer’s instructions. mRNA was transfected in cultured cells to validate activity using a Cel-I enzymatic mutation detection assay. The ZFN mRNA (10ng/μl) was then microinjected into 330 mouse BDF1 embryos and implanted into pseudopregnant female mice recipients.

Genotyping of founders (F_0_) was done using genomic PCR with the primers following primers provided by Sigma, forward 5’-CGCGATGACAGTTTCCAGTA-3’ reverse 5’-CAAACAAAAACCCACTGAGGA-3’. In order to perform DNA sequence analysis, amplified bands of the ZFN target site were cloned in to the pGEM-T Easy Vector system (Promega, Madison, WI) and sequenced by the University of Texas at Austin’s DNA Sequencing Facility. Two founders (line 3 and 11) were then selected for backcrossing and then mated with the wild-type C57BL/6J mice. Speed congenics was performed to select the heterozygous males from each generation for breeding, and to ensure that >99% of the BDF1 background was replaced with the C57BL/6J background. Line 11 was backcrossed a total of 6 generations before proceeding with experiments.

### Diet induced obesity studies

At the age of three weeks, mice (n=8) were placed on high fat chow (60% fat kCal, TD.06414) or control chow (10% fat kCal, TD.0.8806)) for 6 weeks. All diets were provided from Teklad Laboratories (Chicago, IL). At the end of the 6 week treatment period mice were sac’d and mitochondria was extracted from SKM and BAT using the previously described procedure.

### Fasting and cold-exposure studies

Male mice between the ages of 7.5-8.5 weeks old were fasted for 18hrs in individually housed cages with fresh bedding. The body weight of each mouse was recorded before and after fasting treatment. Any mouse under 21g was excluded from the study. Cold tolerance was then tested by placing mice in a temperature-controlled room at 4°C for 3-6 hours, or until body temperature dropped below 25°C. Core body temperatures was recorded prior to cold-exposure and every hour during treatment with a rectal temperature probe (Physitemp, Clifton, NJ).

### Mitochondrial isolation

C2C12 cells were homogenized with a 27.5 gauge needle in CP-1 buffer supplemented with protease and phosphatase inhibitors. Homogenates were then centrifuged at 500g for 10 minutes at 4°C twice, making sure to discard pellets and re-aliquot supernatant in to new tubes in between spins. After second spin, the supernatant was collected and spun at 10,500 g for 10 minutes at 4°C to pellet mitochondria.

Brown adipose tissue, SKM, and heart tissue samples were isolated from mice and finely minced in either CP-1 buffer or 100mM KCl, 500mM Tris-HCl, 2mM EGTA, 1mM ATP, 5mM MgCl2, 0.5% BSA, pH 7.4 supplemented with protease and phosphatase inhibitors. Tissues were then homogenized with a Teflon pestle drill press in smooth glass homogenizers. Homogenates were then centrifuged at 800g for 10 minutes at 4°C twice, making sure to discard pellets and re-aliquot supernatant in to new tubes in between spins. After the second spin, homogenates were applied to a 70μm cell strainer and then centrifuged at 10,500g for 10 minutes at 4°C to pellet mitochondria.

### Immunoblotting

Lysates were prepared in RIPA buffer (50mM Tris-HCl, 1% NP40, 0.5% Sodium Deoxycholate, 0.1% Sodium Dodecyl Sulfate (SDS), 150mM Sodium Chloride (NaCl), 2mM EDTA, pH 8.0) supplemented with protease and phosphatase inhibitor cocktails (Roche, Nutley, NJ). A bicinchoninic acid assay (Pierce Biotechnology, Rockford, IL) was used to quantitate proteins.

Nitrocellulose membranes were probed with the following primary antibodies; rabbit polyclonal α-V5 (Abcam, Cambridge, MA), mouse monoclonal α-V5 (Life Technologies, Grand Island, NY), α-mouse monoclonal α-myc (Cell Signaling, Danvers, MA), rabbit polyclonal α-ECH1 (custom antibody generated by Washington Biotechnology, Columbia, MD), rabbit polyclonal α-ECH1/ECH1 (Abcam, Cambridge, MA), rabbit polyclonal α-UCP3 (custom antibody generated by Washington Biotechnology, Columbia, MD), and rabbit polyclonal anti-UCP3 (Abcam, Cambridge, MA). Following incubation with primary antibodies, the nitrocellulose membranes were probed with α-rabbit-HRP or α-mouse-HRP (GE Healthcare, Piscataway, NJ). Membranes were developed using Super Signal West Pico chemiluminescent (Pierce Biotechnology, Rockford, IL).

### Co-immunoprecipitation

Immunoprecipitation samples (IP) were prepared using 250-500*µ*g of protein and adjusted to 300μl with the previously described immunoprecipitation buffer. Samples were then incubated with either 1μg of primary antibody or IgG (for controls) at 4°C for 16 hours with rotation. Protein G Sepharose beads were blocked in 5% BSA in PBS overnight as previously described. Thirty microliters of Protein G Sepharose beads were then added to each immunoprecipitation sample and rotated at 4°C for 5 hours. Samples were washed 8 times with wash buffer, after then prepared for SDS polyacrylamide gel electrophoresis (PAGE).

### Quantitative RT-PCR

Total RNA was extracted from C2C12 cells and mouse tissues using the TRIzol® reagent (Invitrogen, Carlsbad, CA) according to manufacturer’s instructions. RNA was then reverse transcribed using the TaqMan^®^ Reverse Transciption Kit (Life Technologies, Grand Island, NY) and quantative RT-PCR was performed with the SYBR Green dye using a real-time PCR system (Bio-Rad, Hercules, CA). The following primers used for amplification are listed in appendices. Experiments were repeated in triplicates and data represent fold change relative to levels of GAPDH.

### Immunocytochemistry

Twenty-four hours after transfection C2C12s were fixed and permeabilized in 2.5% paraformaldehyde and 0.1 % TritonX-100 at RT for 20 minutes. The cells were then incubated with either anti-Myc or the peroxisomal marker anti-catalase (Abcam, Cambridge, MA), followed by secondary anti-mouse Alexa 568 or anti-rabbit Alexa 488. All fluorescent images were acquired with a Nikon Eclipse Ti-S microscope, 60x Nikon Plan Apo VC Oil objective with numerical aperture 1.40. Images were captured with Photometrics Coolsnap EZ camera and processed using Nikon NIS Elements BR 3.0 software.

### Myocyte oxygen consumption

Respiration rates in SKM myocytes were quantified using a fiber-optic fluorescence oxygen monitoring system (Instech Laboratories, Plymouth Meeting, PA). Three million cells were added to the 1mL chamber containing Dulbecco’s Modified Eagle Medium from Sigma (St. Louis, MO). Basal and oligomycin-induced uncoupled respiration rates were determined from linear regions of the slope observed after each treatment.

### Fatty acid oxidation assay

C2C12 stable cell lines were grown in a 24-well plate were transiently transfected and assayed for oleate oxidation using the method of Mao et al. (39). Briefly, myocytes were serum starved for 2 hours in substrate-limited media. They were then incubated in 1μC/ml ^14^C-oleic acid (American Radiolabeled Chemicals, St. Louis, MO), lysed with 70% percholoric acid, and collected radioactivity was measured using a scintillation counter.

### Statistics

Analysis of variance comparisons between genotypes were analyzed by a student’s t-test or single factor ANOVA followed by Tukey’s post hoc test, with a p<0.05 set *a priori* as statistically significant.

## Results

### Identification of the interaction between ECH1 and UCP3

UCP3 is a 6-transmembrane (TM) domain-containing protein with 7 hydrophilic (HD) connecting loop domains (Figure 1A, HD1-7). We adapted a yeast two hybrid screen to identify UCP3 binding partners and used the seven HD UCP3 domains as baits in yeast expressing a human heart cDNA library, leading to the identification of ECH1 as a candidate UCP3 interacting protein. This approach revealed that HD1, 3, and 4, but not HD2, 6, or 7 interacted with full length ECH1 (Fig. 1B).

**Figure 1.**
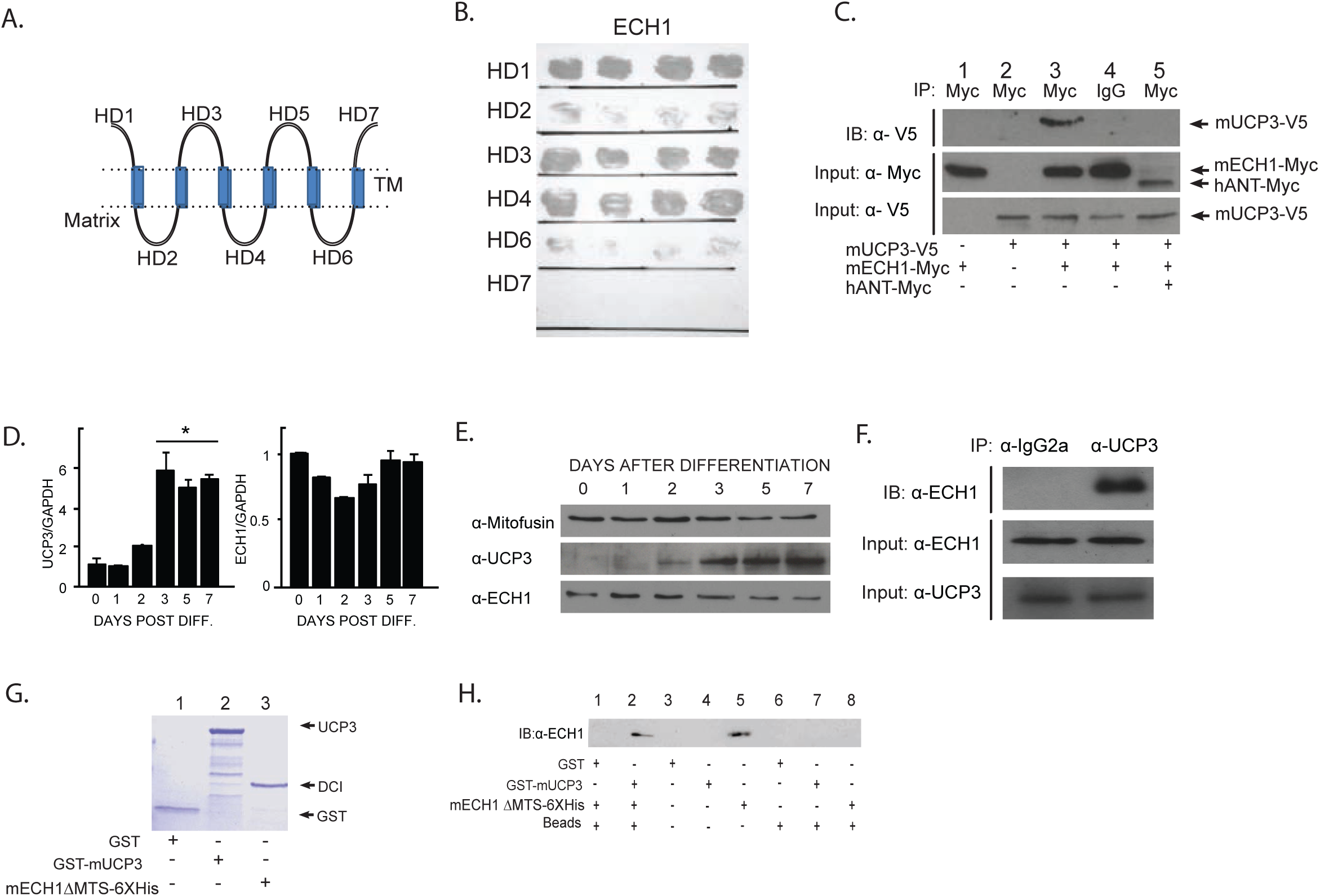
Interaction of ECH1 and UCP3. (A) UCP3 is expressed on the inner mitochondrial membrane and contains seven hydrophilic domains (HD1-7) connected by 6 transmembrane domains. (B) Growth phenotypes of yeast expressed full-length ECH1 and individuals UCP3 domains HD1-7. (C) Co-immunoprecipitation of ECH1-Myc and UCP3-V5 in co-transfected cells. Cells were transfected with the indicated plasmids. Immunoprection (IP) was performed with anti-Myc or negative control IgG. Co-precipitating proteins were detected by immunoblotting with anti-V5. Input expression levels were confirmed in lower panels. (D-E) Endogenous mRNA and protein expression profiles of ECH1 and UCP3 in differentiating mouse myoblasts C2C12s for 1-7 days (n=3). (F) Co-immunoprecipitation of endogenous UCP3 and ECH1 in differentiated C2C12s. Mitochondrial lysates isolated from myotubes were immunoprecipitated (IP) with anti-UCP3 or IgG. Co-precipitating proteins were detected by immunoblotting with anti-ECH1. Endogenous protein levels of UCP3 and ECH1 were confirmed as indicated. (G) Expression of purified recombinant GST-UCP3 (lanes 2), ECH1ΔMTS-6XHis (lanes 3), and GST vector control (lane 1) from BL21 (DE3) E. Coli. (H) UCP3 and ECH1 bind directly in vitro. Bacteria were transformed with the indicated plasmids and lysates were incubated with glutathione-coupled sepharose beads (Beads). *In vitro* translated ECH1 was immunoprecipitated only with GST-mUCP3 alone (lane 2). Lysates of cells expressing ECH1ΔMTS-6XHis alone (lane 5) served as a positive control for *in vitro* translation.

We first tested the ECH1:UCP3 interaction in mammalian cells by co-transfecting cells with a V5-tagged mouse UCP3 construct (UCP3-V5) and Myc-tagged mouse ECH1 construct (ECH1-Myc). Lysates were immunoprecipitated with anti-Myc antibody and co-immunoprecipitated UCP3-V5 was detected by immunoblotting with anti-V5 antibody. Results showed that UCP3-V5 was immunoprecipitated with anti-Myc (Fig. 1C, lane 3), but not by IgG (lane 4). Furthermore, UCP3-V5 was not immunoprecipitated with the Myc-tagged human adenine nucleotide translocase (hANT-myc, Fig. 1C, lane 5), thus confirming the specificity of the mitochondrial ECH1:UCP3 interaction. To rule out overexpression artifacts, we next tested whether complex formation between ECH1 and UCP3 was detectable at endogenous levels. Consistent with previous findings (Nagase et al. 2001; Solanes et al. 2000; Son et al. 2001), C2C12 mouse myoblasts that were differentiated in low serum media (2% equine serum) showed an induction of UCP3 mRNA and protein expression levels (Fig. 1E, upper blot, days 2-6). Interestingly, the data showed that endogenous ECH1 mRNA and protein levels did not change upon differentiation (Fig. 1D). We then tested whether the ECH1:UCP3 complex could be detected in differentiated C2C12 myotubes when both proteins were present at physiological concentrations. Using lysates from differentiated myotubes, we found that endogenous ECH1 could be detected when immunoprecipitated with anti-UCP3, but not with rabbit anti-IgG in C2C12 myotubes (Fig. 1F).

To test whether UCP3 and ECH1 formed a direct complex, we used purified recombinant UCP3 and ECH1 proteins and performed *in vitro* pull-down experiments. Full length UCP3 protein with an N-terminal GST tag (GST-UCP3) was expressed and purified from bacteria, along with recombinant ECH1 protein that lacked its mitochondrial targeting sequence and was linked to a C-terminal 6X His tag (ECH1ΔMTS-6XHis) (Fig. 1G). Immunoprecipitation experiments with the purified proteins demonstrated that GST-UCP3 was able to pull down ECH1ΔMTS-6XHis *in vitro* (Fig. 1H, Lane 2). As expected, ECH1ΔMTS-6XHis was not pulled down in either beads (Lane 8) or GST (Lane 1) alone.

### Endogenous ECH1 protein expression and localization

The endogenous function(s) of ECH1 in a physiological context are unclear. Interestingly, ECH1 exhibited an overlapping protein expression profile with UCP3 in highly metabolic tissues including brown adipose (BAT), skeletal muscle (SKM), and heart (HRT) (Fig. 2A). ECH1 is unique in that it contains both mitochondrial (MTS) and peroxisomal targeting sequences (PTS) that flank a catalytic and trimerization domain (Modis et al. 1998; Zhang et al. 2001), but its sub-cellular distribution in muscle (or tissues other than liver? Or any tissue?) has not been described. To examine whether ECH1 localization is predominantly mitochondrial or peroxisomal, immunocytochemistry was performed in C2C12 myocytes that were co-transfected with ECH1-Myc and either Mito-GFP or empty vector control plasmid. Pearson’s correlation coefficient was used to quantify the subcellular localization of ECH1, and revealed that ECH1 localizes predominantly to mitochondria based on the co-localization of immunofluorescent staining for ECH1 with Mito-GFP (Fig. 2C). Endogenous staining of catalase was used as a peroxisomal marker.

**Figure 2.**
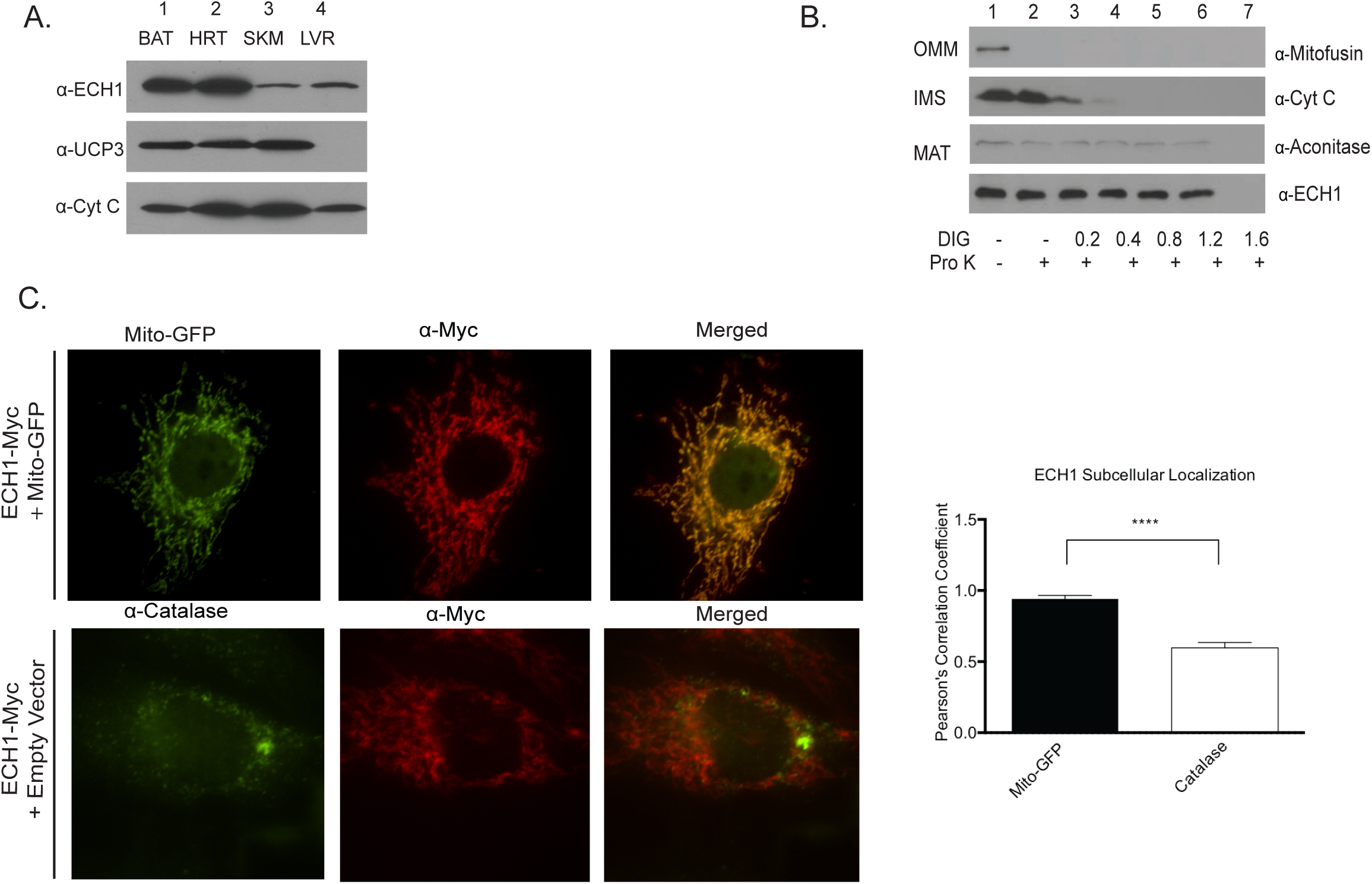
Characterizing ECH1 and UCP3 expression and localization. (A) Western blots of ECH1 and UCP3 in mitochondrial lysates isolated brown adipose tissue (BAT), skeletal muscle (SKM), heart (HRT), and liver (LVR) tissues (n=4). (B) Sub-mitochondrial localization assay demonstrates ECH1 localizes to the mitochondrial. Mitochondria isolated from C2C12 myotubes were treated with proteinase K (Pro K) (lanes 2-7) and increasing concentrations of the detergent digitonin (DIG) (lanes 3-7, 0.2 to 1.6 mM DIG). Immunoblots detected presence of the outer mitochondrial membrane (OMM) protein mitofusin, mitochondrial membrane space (IMS) resident protein cytochrome c (Cyt C), and the matrix (Mat) resident protein aconitase. ECH1 protein expression was similar to aconitase, thus indicating matrix localization. (C) ECH1 sub-cellular localization. Representative immunocytochemistry images of C2C12 myoblasts co-transfected with ECH1-Myc and either the GFP fusion protein containing a mitochondrial targeting sequence (Mito-GFP, top panel) or empty vector control (bottom panel). Cells were incubated with primary antibodies anti-Myc (ECH1-Myc, top and bottom panel) and anti-catalase (peroxisome marker, bottom panel), followed by incubation with corresponding fluorescent secondary antibodies anti-mouse (red, mDCI-Myc) and anti-rabbit (green, bottom panel, catalase). (D) Pearson’s correlation coefficient for co-localization of ECH1-Myc (red) and Mito-GFP or catalase (green). Data are expressed as means ± SEM from three independent experiments (n=6-7 cells) ***p<0.001, statistical significance was detected by Student’s t test.

It is likely that UCP3 shares a similar protein structure to other members of the mitochondrial carrier protein family such as ANT. Based on the topology predictions and crystal structure of ANT (Notario et al. 2003), it is thought that these proteins have N- and C-terminal hydrophilic domains that face both sides of the inner mitochondrial membrane. Thus, the hydrophilic UCP3 domains 1, 3, 5, and 7 are localized in the intermembrane space, and domains 2, 4, and 6 localize to the matrix. Because the bait constructs HD1, 3, & 4 captured ECH1 in the yeast two-hybrid analysis (Fig. 1 A-B) it is possible that ECH1 could interact with UCP3 on either or both sides of the inner mitochondrial membrane. In order to define the sub-mitochondrial localization of ECH1 we performed a digitonin-based localization assay on mitochondria isolated from C2C12 myotubes. As indicated in Fig. 2B, mitochondria were treated with proteinase K (Pro K) and increasing amounts of the membrane permeabilizing detergent digitonin (DIG). Fig. 2B immunoblots show that in the absence of DIG and Pro K treatment, the resident mitochondrial proteins of the outer mitochondrial membrane (OMM, mitofusin), intermembrane space (IMS, Cyt C), and matrix (MAT, aconitase) could all be detected (Fig. 2B, lane 1). Treatment of mitochondria with Pro K alone (lane 2) resulted in the selective digestion of the OMM protein mitofusin. In the presence of Pro K and increasing concentrations of DIG (lanes 3-7, 0.2 to 1.6 mM DIG), IMS Cyt C immunoreactivity was lost at lower DIG levels, whereas the matrix (MAT) resident aconitase required increased digitonin concentrations for proteolysis. ECH1 was protected from the Pro K and DIG treatments to a similar degree as aconitase, indicating that ECH1 localizes to the mitochondrial matrix.

### ECH1 interacts with the central matrix loop of UCP3

As mentioned above, the yeast two hybrid results identified HD 1, 3, and 4 as potential interaction motifs. Next, we sought to map the UCP3-interaction motifs that are necessary for complex formation with ECH1 by generating a series of V5-tagged (C terminus) UCP3 truncation mutants lacking the indicated HD displayed in Figure 3A. Cells co-transfected with ECH1-Myc and either full-length UCP3-V5 or the UCP3 truncation mutants were lysed and immunoprecipitated with anti-Myc. Absence of hydrophilic domain 4 (ΔHD4) caused a significant decrease in the amount of UCP3 co-immunoprecipitated with ECH1-Myc (Fig. 2D, lane 7). These results indicate that complex formation between ECH1 and UCP3 is mediated through HD4, which is the central matrix loop of UCP3. Furthermore, additional co-immunoprecipitation studies with V5-tagged HD4 mutants, each lacking one third of the domain, showed that ECH1 and UCP3 likely interact via the first 12 amino acids of the central matrix loop of UCP3 (Figure S1).

**Figure 3.**
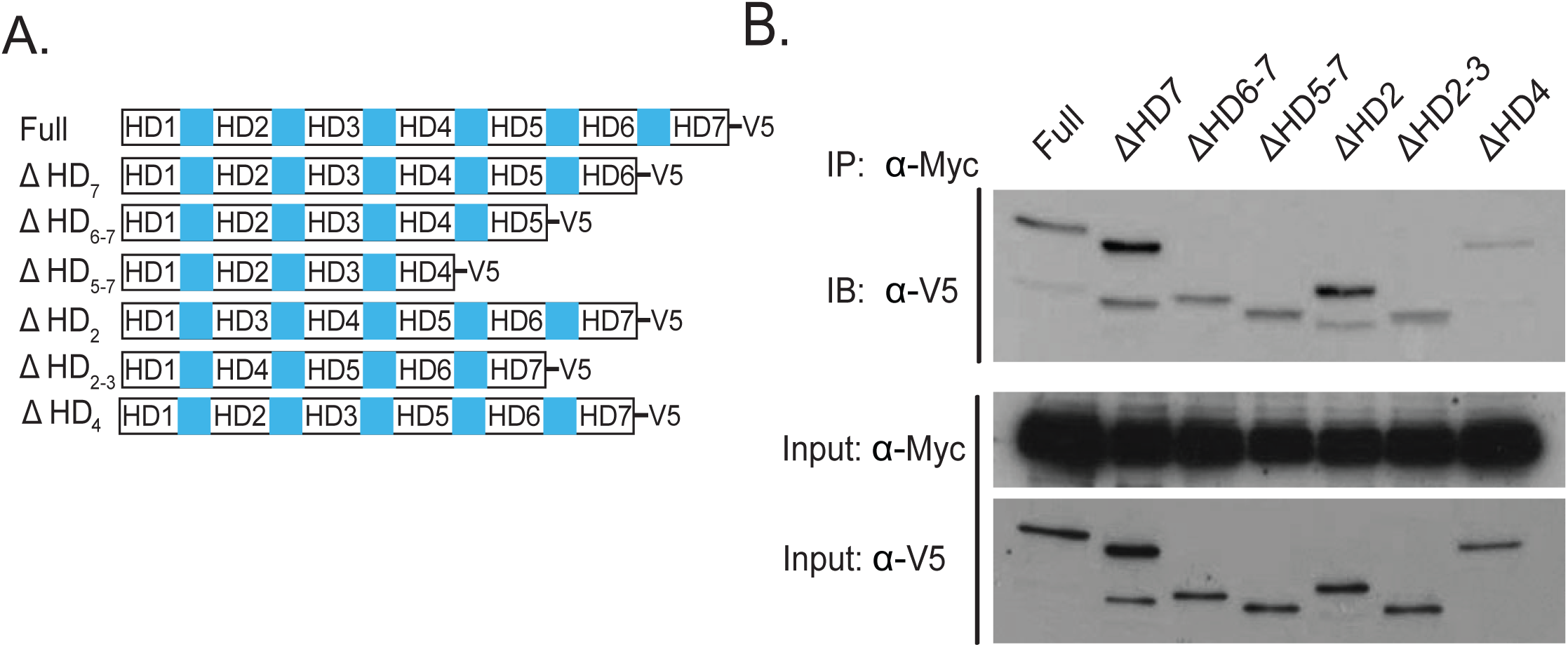
Mapping the ECH1:UCP3 interacting domains. (A) Depiction of the mUCP3-V5 hydrophilic domain (HD) truncation mutant constructs used to identify ECH1 interacting motifs. (B) ECH1 interacts with HD4 of UCP3. Cells were co-transfected with ECH1-myc and the indicated V5 tagged full length UCP3 or UCP3 HD mutants as indicated (top). Lysates were immunoprecipitated (IP) with anti-Myc antibody and immunoblots (IB) were probed with anti-V5. Middle panel shows input for ECH1-myc, and lower panel shows input for UCP3 and UCP3 HD mutants.

### Fatty acid regulation of complex formation between ECH1 and UCP3

We then focused on defining the biochemical and physiological factors that regulate ECH1 and UCP3 complex formation. We generated Myc-tagged ECH1 catalytic mutants to address whether the mutants could be co-immunoprecipitated with UCP3-V5 in lysates extracted from co-transfected cells (Fig. 4A). According to previous structural and mechanistic studies of ECH1, the Aspartic acid 204 (D204) and glutamic acid 196 (E196) residues located in the active site of ECH1 are essential for catalysis (Modis et al. 1998). Aspartic acid 176 (D176) is also located in the active site of ECH1, but is shown to have little effect on the enzyme’s catalytic activity (Zhang et al. 2001). Co-immunoprecipitation of UCP3-V5 with both Myc-tagged ECH1 catalytic mutants D204N-Myc and E196-Myc was severely diminished compared to the control, ECH1I-Myc (Figure 4A). Interestingly, the ECH1 mutant D176N-Myc that still possesses catalytic activity did not have the same attenuated effect on complex formation with UCP3-V5. However, introducing the E196Q mutation to generate D176N/E196Q-Myc restored the loss of interaction with UCP3-V5, thus indicating that ECH1 catalytic activity is important in ECH1 and UCP3 complex formation (Fig. 4A).

**Figure 4.**
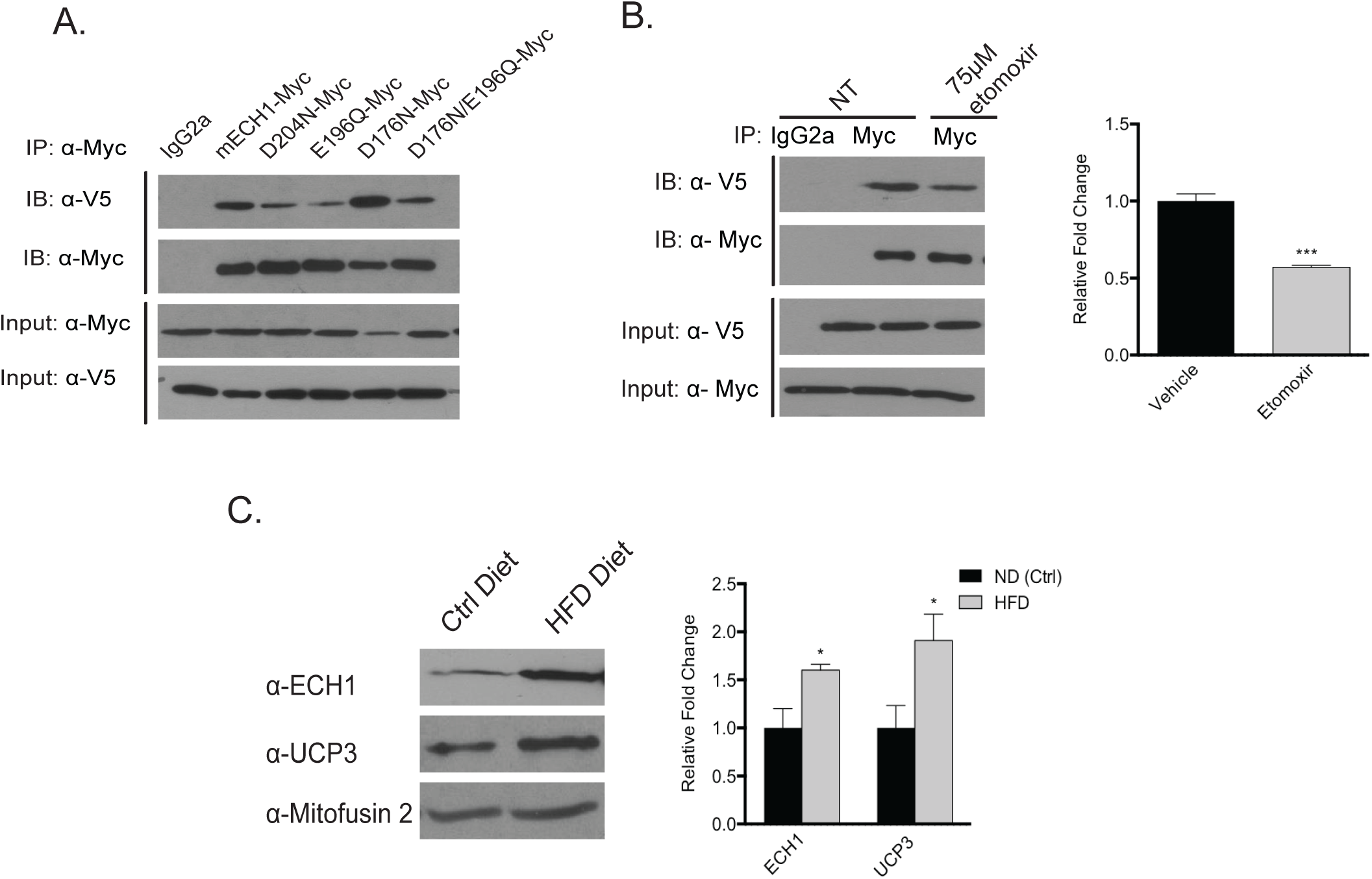
Fatty acid regulation of the ECH1 and UCP3 interaction. (A) Loss of ECH1 catalytic activity diminishes complex formation between ECH1:UCP3. Lysates were immunoprecipitated with either IgG or anti-Myc from cells co-transfected with UCP3-V5 and either ECH1-Myc or the Myc-tagged ECH1 catalytic mutant constructs (D204N-Myc and E196Q-Myc, lanes 1-3). To rule out any effects independent of catalytic activity we generated an ECH1 mutant that has been previously shown to retain enzymatic activity (D176N-Myc, lane 4) and the ECH1 double mutant that re-introduced the E196Q catalytic mutation (D176N/E196Q-Myc). Co-immunoprecipitation of UCP3-V5 was detected by immunoblotting with anti-V5 (first panel). Successful immunoprecipitation and pull down of Myc tagged ECH1 and ECH1 mutants were confirmed by immunoblotting with anti-Myc (second panel). Protein expression levels were confirmed in lower panels. (B) Inhibition of fatty acid transport in to mitochondria augments complex formation. Cells co-transfected with UCP3-V5 and ECH1-Myc constructs were treated for 18hrs with either vehicle (lane 1-2) or 75 μM etomoxir, the CPT1 inhibitor. Co-immunoprecipitations were performed as previously described. Densitometry was performed on immunoblots of precipitated UCP3-V5 (top panel) and normalized to pull down of ECH1-Myc (second panel). Data represent means ± SEM from 3 independent experiments. ***p< 0.001, statistical significance was detected by Student’s t test. (C) Protein expression levels of UCP3 and ECH1 are both elevated in skeletal muscle of wild type mice fed high fat diet (60% kCal) for 6 weeks. In densitometry analyses of immunoblots, ECH1 and UCP3 protein expression was normalized to mitofusin-2. Data are expressed as means ± SEM (n=8) *p< 0.05, statistical significance was detected by Student’s t test.

Having established that modifying ECH1 catalytic activity can affect complex formation between ECH1 and UCP3, we reasoned that complex formation between ECH1 and UCP3 might be stimulated by the presence of FAs. To test this hypothesis, we blocked FA import into the mitochondria by inhibiting the key mitochondrial FA transporter, carnitine palmitoyl transferase I (CPT1) with the drug etomoxir. Co-immunoprecipitation experiments were performed in lysates extracted from HEK293T cells co-transfected with UCP3-V5 and ECH1-Myc, and then treated with 75μM etomoxir. Densitometry analyses of immunoblots detecting the amount of UCP3-V5 co-immunoprecipitated with ECH1-Myc showed a significant decrease in ECH1:UCP3 complex formation in the etomoxir treated cells compared to vehicle treated (Fig. 4B). These results indicate that complex formation between ECH1 and UCP3 is stimulated by FAs, and could thus be physiologically relevant in metabolic pathologies where FAs serve as a primary fuel source for mitochondria. Consistent with these findings, we observed a ∼1.5 and 2 fold increase in ECH1 and UCP3 protein expression levels, respectively, in isolated SKM mitochondria from mice fed high-fat diets for 6 weeks (Fig. 4C). Body weights were significantly higher in the mice fed high fat diets, thus affirming that the changes in ECH1 and UCP3 protein expression were correlated with the degree of obesity in our animals (Supplementary Fig. 2).

To characterize the functional influence of the ECH1:UCP3 complex on mitochondrial metabolism in SKM we utilized the Precision Lenti-ORF plasmids (pLOC) to generate stable cell lines in C2C12 myoblasts that either overexpresses ECH1 (C2C12-ECH1) or empty vector pLOC (C2C12-EV). Stables colonies of both cell lines were selected and densitometry analyses confirmed protein overexpression of ECH1 in the C2C12-ECH1 cell lines at physiological levels (Fig. 5A) that were similar to the induction of ECH1 expression in SKM of mice fed high fat diets (Fig. 4C). Given that UCP3 function is closely tied to FA metabolism, if ECH1 forms a direct complex with UCP3 in the presence of FAs it is possible that complex formation could regulate UCP3 activity, thus enhance uncoupled respiration. To investigate this notion we transfected UCP3-V5 in to our stable cell lines C2C12-ECH1 and -EV and measured uncoupled respiration using a fiber-optic fluorescence oxygen monitoring system. As shown in Figure, oligomycin-induced uncoupled respiration was higher in cells overexpressing ECH1 and UCP3-V5 compared to cells expressing UCP3-V5 alone. We then set out to examine the consequences of the ECH1:UCP3 complex formation on the ECH1-dependent metabolism of unsaturated fatty acids through quantification of radiolabeled oleate oxidation in our stable cell lines transfected with UCP3-V5. Consistent with previous observations, UCP3 expression alone led to a slight increase in oleate metabolism (Fig. 5C). Interestingly, ECH1 and UCP3 overexpression led to a synergistic increase in oleate oxidation.

**Figure 5.**
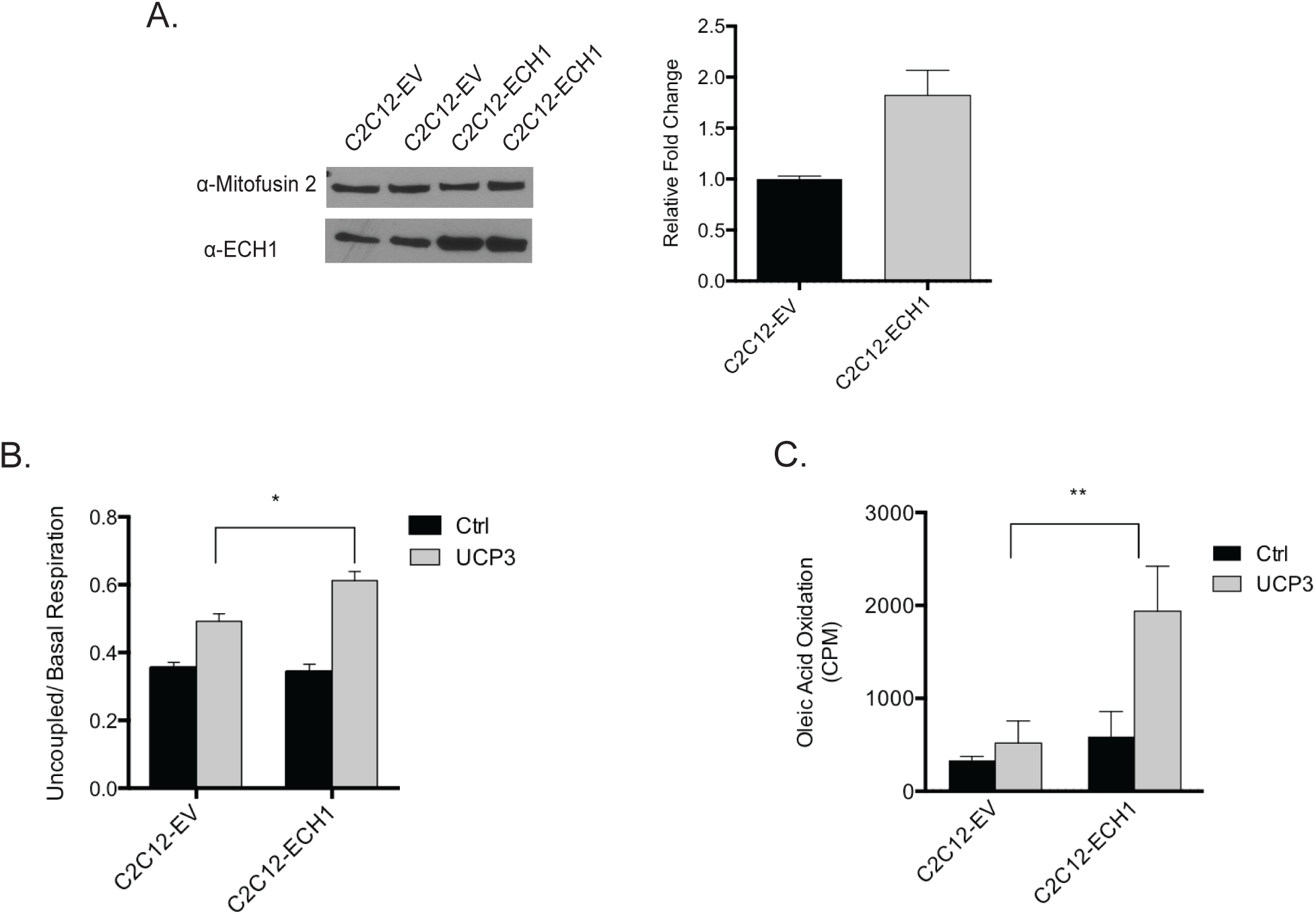
Functional implications of ECH1 and UCP3 in mitochondrial metabolism. (A) Lentiviral overexpression of ECH1 in C2C12. For lentiviral transduction, C2C12s were incubated with lentiviral particles containing either the empty vector Precision Lenti-ORF plasmid (pLOC) or pLOC-ECH1 construct. Two stable colonies of C2C12s expressing empty vector pLOC (C2C12-EV) or pLOC-ECH1 (C2C12-ECH1) were selected with blasticidin and grown up. Densitometry was performed on immunoblots of ECH1 protein expression normalized to mitofusin. (B) ECH1 expression enhances uncoupled respiration. C2C12-EV and C2C12-DCI cells were transfected with UCP3-V5 (UCP3) or empty vector (Ctrl). Uncoupled respiration rates were normalized to basal respiration rates. (C) Fatty acid oxidation was determined by measuring captured ^14^CO_2_ produced from [1-^14^C] oleic acid incubated with transfected C2C12-EV and C2C12-DCI cells. Data representative of two stable colonies and expressed as means ± SEM. *p< 0.05, **p< 0.01, statistical significance was detected by one way analysis of variance (ANOVA) followed by Tukey’s *post hoc* test.

### Physiological Relevance of ECH1 and UCP3 in Metabolic Stress

The thermogenic capabilities of UCP3 remain controversial. Even though UCP3 knockout (UCP3^-/-^) mice do not exhibit a clear cold-intolerant or obese phenotype, there is significant evidence to support the role of UCP3 in maintaining energy balance in metabolically challenging conditions where FA oxidation is high. Indeed, experiments demonstrating that fasted UCP3^-/-^ mice have impaired rates of FA oxidation along with elevated levels of FA matrix accumulation, suggest that UCP3 is necessary for mitochondrial adaptation to fasting (Seifert et al. 2008). Given that FA metabolism and transport is essential in driving cold-induced thermogenesis, we tested whether the absence of UCP3 would affect cold-induced thermogenesis in fasted mice. Suprisingly, we found that UCP3^-/-^ mice (between the ages of 7-8.5 wks) fasted for 18 hours were more sensitive to cold after 6 hours compared to the wild type (Fig. 6A).

**Figure 6.**
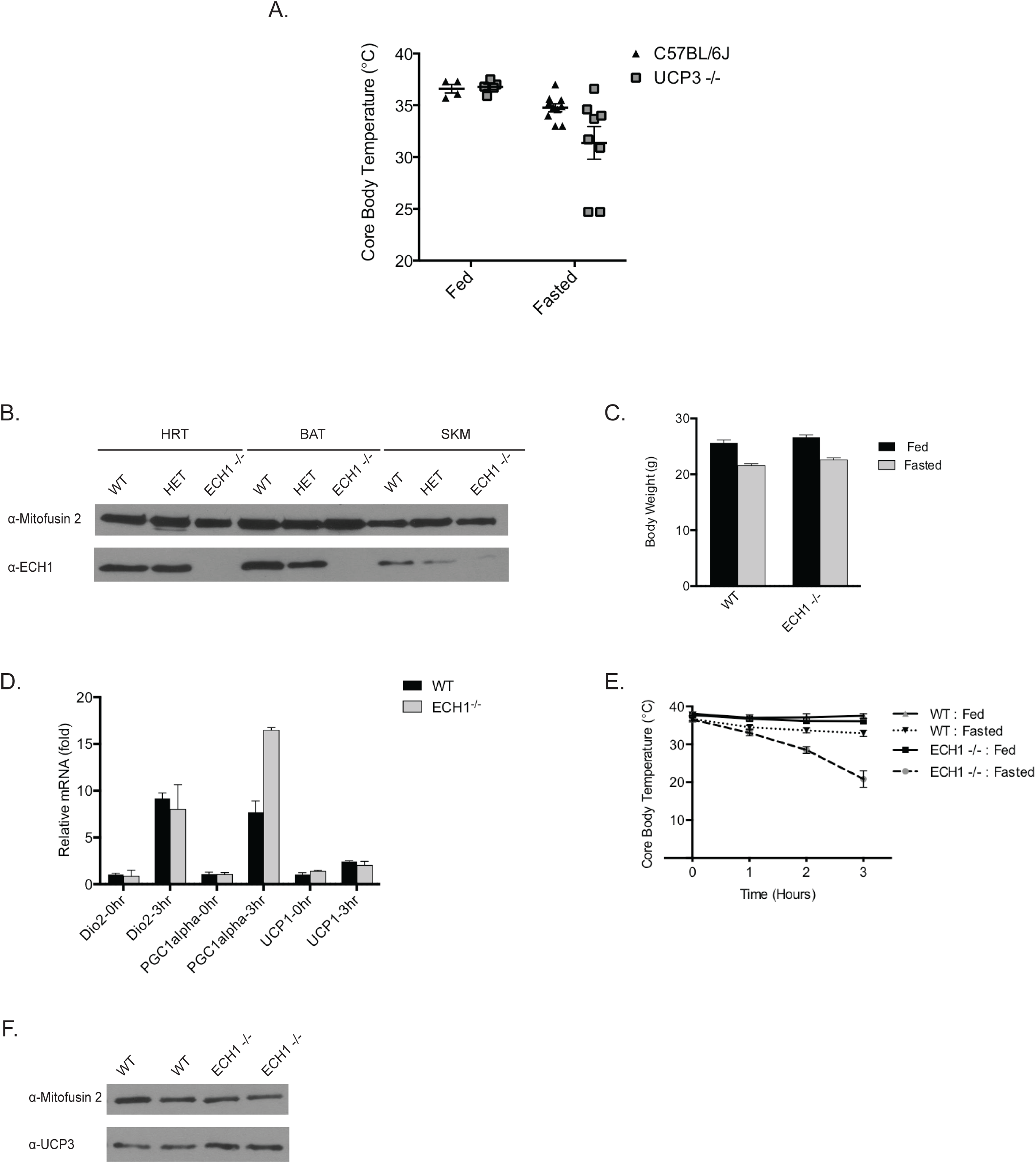
Physiological relevance of ECH1:UCP3 complex in metabolic stress. (A) Body temperature of wild type (WT) and UCP3^-/-^ mice following 18hr fast and 6hr cold challenge 4°C (n=4-10). Error bars represent SEM. *p< 0.05, statistical significance was detected by student’s t test. (B) ECH1 expression in oxidative tissues from WT, HET, and KO (C) Body weights of WT and ECH1^-/-^ before (fed) and after 18hr fast (fasted) (n=4-5). *p< 0.05, statistical significance was detected by student’s t test (D) mRNA expression levels of canonical cold-induced genes (n=4-5). (E) Body temperatures of WT and ECH1 ^-/-^ mice following 18hr fast and 3hr cold challenge at 4°C (F) Expression of UCP3 in skeletal muscle of ECH1KO mice. * p< 0.05, statistical significance was was detected by two way analysis of variance (ANOVA) followed by Tukey’s *post hoc* test. (C-D) Data are expressed as means ± SEM.

Like UCP3, it has been proposed that ECH1 may play a crucial role in the protection against lipid overload in conditions of high metabolic stress when FAs serves as the primary source of energy. Indeed, the most common phenotype seen with whole body knockout mouse models of different auxiliary unsaturated FA metabolizing enzymes is severe cold intolerance with prior fasting (Miinalainen et al. 2009; Janssen & Stoffel 2002). In order to characterize the extent to which ECH1 contributes to non-shivering thermogenesis in periods of severe metabolic stress we generated a global ECH1 knockout mouse model (ECH1^-/-^) utilizing a zinc finger nuclease based-targeted genomic approach. The custom made ECH1-ZFNs, comprised of a DNA binding domain targeted to exon 3 of the mouse ECH1 gene located on chromosome 7, was fused to a Fok I nuclease domain. Genomic PCR and sequence analysis was used to check ZFN-mediated mutations (Supplemental Fig.2A-C).

Analyses of ECH1 expression by western blot confirmed a complete loss of ECH1 expression in SKM, BAT, and HRT (Fig. 6B). We then subjected these mice to 18hr fasting and cold stress (4°C) and monitored changes in core body temperature. Similar to the wild-type group, the ECH1-/- mice showed a slight decrease in body temperature following fasting treatment compared to the fed groups (Fig. 6C, 0 hrs). However, the fasted ECH1 -/-mice were unable to maintain core body temperature during acute cold exposure at 4°C (Fig. 6D). Interestingly, the body weights of the wild-type and ECH1^-/-^ mice were not different in the fasted and fed groups, thus indicating that the thermogenic phenotype seen in the ECH1^-/-^ mice was independent of any differences in body weight following fasting (Fig. 6E). We also found that the ECH1^-/-^ mice did not exhibit a difference in mRNA expression levels of the canonical cold-induced thermogenic genes in brown adipose tissue UCP1, Dio2, and Pgc1 (Fig. 6F). These results suggest that the signaling downstream of adrenergic stimulation of BAT in response to cold exposure is intact in the ECH1-/-mice and is not likely contributing to the severe cold-intolerant phenotype. Surprisingly, when we looked at UCP3 protein expression in SKM following fasting and acute cold exposure treatments, we found that ECH1-/-mice had higher levels of UCP3 compared to wild type. Taken together, this compensatory increase in UCP3 corroborates the synergistic relationship between ECH1 and UCP3 in skeletal muscle metabolism. Thus, the ECH1-/-mice are unable to maintain core body temperature despite the compensatory increase in UCP3 protein expression in SKM.

## Discussion

There is considerable evidence to support that UCP3 has the ability to regulate FA metabolism and transport to protect SKM mitochondria against FA overload (lipotoxicitiy) and insulin resistance, but only when activated in certain physiological contexts. FAs have been implicated as potent activators of UCP function (Jiménez-Jiménez et al. 2006; Skulachev 1999; Hagen & Lowell 2000), however significant debate exists regarding the mechanisms by which FAs activate UCP3, and by which UCP3 regulates FA metabolism. Our work demonstrating that the auxiliary unsaturated FA metabolizing enzyme ECH1 interacts with UCP3 in the mitochondrial matrix, at endogenous levels, supports a mechanism by which UCP3 can facilitate FA transport and influence mitochondrial metabolism. Importantly, we also demonstrate that UCP3 and ECH1 are both important in facilitating an adaptive response to metabolic stress, and that complex formation between these proteins could be key in regulating a thermogenic response to acute cold exposure in a fasted state.

The characterization of the ECH1:UCP3 complex sheds new light on how the specific metabolism of polyunsaturated FAs with odd-numbered double bonds contributes to mitochondrial energy balance. ECH1 is a unique enzyme in that it demonstrates dual-organelle distribution to the peroxisomes and mitochondria in mammalian cells. The enzyme contains a known type I peroxisomal targeting sequence (SKL) in its C-terminus (Filppula 1998), and it has also been shown that the first 40 amino acids in the N terminus resembles a cleavable mitochondrial targeting signal as predicted by Mitoprot II analysis (Claros & Vincens 1996). While mitochondrial FA metabolism is a major contributor to energy production and balance, peroxisomal FA oxidation is not. Indeed, FA metabolism in peroxisomes is only capable of shortening the fatty acyl-CoA chains, while the complete degradation of the fatty acyl-CoA chain to generate ATP occurs exclusively in the mitochondria. Therefore, our finding that ECH1 localizes primarily to the mitochondria rather than peroxisomes is consistent with previous studies that ECH1 is important in maintaining pools of coenzyme A and regulating energy balance.

It has been proposed that ECH1 plays an important role in the disposal of unmetabolizable FA metabolites (Shoukry & Schulz 1998). ECH1 is an essential auxiliary enzyme in the reductase-dependent pathway that is responsible for catalyzing the isomerization of the 3,5-dienoyl-CoA substrate to 2,4-dienoyl-CoA. This is regarded as a crucial step in the reductase-dependent pathway because unlike other FA-intermediates that can be metabolized by redundant FA enzymes, the 3,5-dienoyl-CoA substrate is a “dead end metabolite” that is exclusively metabolized by ECH1. Based on these realizations it was later suggested that the reductase-dependent pathway be renamed to the ECH1-dependent pathway (Shoukry & Schulz 1998) given that in the absence of ECH1 activity, the “dead end metabolite” would accumulate, sequester coenzyme A, and halt mitochondrial β-oxidation.

Interestingly, the proposed physiological function of ECH1 is similar to previous theories by Harper et al. that suggest UCP3 also protects against lipotoxicity by facilitating FA transport to maintain coenzyme A availability in conditions that require high levels of FA oxidation. In support of this notion, UCP3 expression is induced in response to high fat feeding and fasting, thus suggesting a role for UCP3 in conditions where the rates of mitochondrial FA oxidation are high. Furthermore, it has also been shown that UCP3 overexpression in SKM can lower circulating levels of acylcarnitines and thus facilitate complete FA oxidation (Aguer et al. 2013). Based on these findings, it makes sense that UCP3 would interact with a FA metabolizing enzyme to mediate efficient FA oxidation. Accordingly, our data demonstrating that ECH1:UCP3 complex formation is regulated through FAs and likely activated in conditions that require enhanced levels of FA oxidation e.g. high fat feeding, suggests these two proteins could be facilitating a compensatory response to metabolic FA overload. Indeed, our data proposes a modified model by which UCP3 functions to maintain coenzyme A availability in part through ECH1-dependent metabolism of unsaturated FAs. The finding that UCP3 and ECH1 expression synergistically increases oleic acid oxidation suggests that UCP3 binding could enhance ECH1 activity and promote unsaturated FA oxidation. In addition to this, we also demonstrate that ECH1 and UCP3 expression synergistically enhances uncoupled respiration in C2C12 myocytes. Taken together, it is tempting to speculate that complex formation between ECH1 and UCP3 functions to coordinate FA oxidation and uncoupling activity in a compensatory pathway that is activated in situations that require enhanced levels of FA oxidation.

Despite the numerous debates regarding the thermogenic capabilities of UCP3 in SKM, results herein demonstrate the UCP3 is important in mediating an adaptive thermogenic response to cold temperatures, in a fasted state. The assumption that UCP3 is not a physiological thermogenic regulator stems from the finding that UCP3^-/-^ mice are able to maintain core body temperature in response to cold. However, the lack of a cold-phenotype may be due to a compensatory process, but it does not necessarily exclude the possibility that UCP3 can contribute to whole-body thermogenesis through alternative mechanisms that may be crucial in different physiological contexts. Indeed, UCP3 has been shown to be a crucial molecular mediator of non-shivering thermogenesis in response pharmacological amphetamines (Mills et al. 2003), and thyroid hormone (Flandin et al. 2009) which is arguably one of the most important hormonal regulators of whole body thermogenesis and energy metabolism. Indeed, observed changes in UCP3 expression in certain physiological contexts where mitochondria rely predominantly on FA metabolism, along with mechanistic studies in fasted mice, suggest that UCP3 may be relevant in mediating an adaptive increase in FA oxidation capacity in SKM (Seifert et al. 2008). Given that fasting and cold exposure are both stimulators of FA oxidation, it is likely that the absence of UCP3 diminishes efficient FA handling in response to these metabolic stressors, thus diminishing optimal cold-induced thermogenesis in UCP3^-/-^ in a fasted stated.

Lastly, we demonstrate for the first time that unsaturated FA metabolism through the reductase/ECH1-dependent pathway is essential for thermogenesis in conditions of severe metabolic stress. The impaired thermogenic phenotype seen in the ECH1^-/-^ mice in response to fasting is similar to previous studies with mice lacking dienoyl-CoA reductase, an auxiliary enzyme involved in the metabolism of all species of unsaturated FAs (odd- and even-numbered double bonds)(Miinalainen et al. 2009). Despite previous work indicating that the reductase/ECH1-dependent pathway contributes to a minor portion of mitochondrial β-oxidation of unsaturated FAs with odd-numbered double bonds (Shoukry & Schulz 1998), our work clearly shows that impairment of this metabolic pathway can have a significant effect on whole-body thermogenesis in fasted conditions. Taken together, this data shows that ECH1 is an essential regulator of cold-induced thermogenesis in a fasted state, through a mechanism that is mediated in part through UCP3-dependent activity in SKM.

## Abbreviations used

(UCP): Uncoupling proteins
(UCP3): Uncoupling protein 3
(SKM): Skeletal Muscle
(NST): non-shivering thermogenesis
(FA): Fatty acid
(ECH1) BAT, HD, ANT, TM, MTS, DIG, MAT, IMS, OMM: Δ^3,5^Δ^2,4^dienoyl-CoA isomerase

